# Human XIRP1 is a macrophage podosome protein utilized by *Listeria* for actin-based motility

**DOI:** 10.1101/2022.08.28.505595

**Authors:** Rodolfo Urbano, Eui-Soon Park, Kyle Tretina, Alexandru Tunaru, Ryan G. Gaudet, Xiaoyun Hu, Da-Zhi Wang, John D. MacMicking

**Affiliations:** Howard Hughes Medical Institute; Yale Systems Biology Institute; Department of Microbial Pathogenesis Yale University School of Medicine, New Haven, United States; Department of Immunobiology, Yale University School of Medicine, New Haven, United States; Department of Cardiology, Boston Children’s Hospital, Harvard Medical School, Boston, United States

## Abstract

Actin is integral to eukaryotic physiology as a biomechanical polymer and as a structural barrier for cell-autonomous defense against infection. Some microbial pathogens exploit the actin cytoskeleton, however, to evade cell-autonomous immunity. Subversion of actin to enter host cells and for actin-based motility are often employed by intracellular pathogens to spread from cell-to-cell. Using RNA-sequencing and computational data mining, we identify the host actin-binding protein XIRP1 as commonly induced during infection. XIRP1 is expressed by fibroblasts and macrophages in response to immune cytokines such as interferon-gamma (IFN-γ) and infection with bacteria such as *Listeria, Shigella,* and *Salmonella*. Confocal and super-resolution structured illumination microscopy (SIM) found XIRP1 localizes to fibroblast focal adhesions and macrophages podosomes. Within human macrophages, XIRP1 is recruited to cytosolic *Listeria monocytogenes* in an ActA-dependent manner as it replicates and uses actin-based motility for host cell escape. Chromosomal removal of XIRP1 in mice impaired this dissemination and rendered them more resistant to *Listeria* infection than C57BL/6NJ wildtype controls *in vivo*. We propose that professional cytosolic pathogens like *Listeria* can co-opt XIRP1 to escape the hostile intracellular environment of IFN-γ-activated macrophages as part of the host-pathogen arms race during cell-autonomous immunity.

## Introduction

Cell-autonomous immunity is an ancient defense strategy deployed by all living organisms to combat infection (MacMicking, 2012; Randow et al., 2013; Wein & Sorek, 2022). In eukaryotes, this defensive strategy extends to preventing pathogen escape and cell-to-cell spread (Mostowy & Shenoy, 2015). Host actin is central to these interactions and underlies both protective immunity and microbial mechanisms of pathogenesis. As one of the main structural components of eukaryotic cells, the actin cytoskeleton consists of actin and associated actin-binding proteins which function as scaffolds and regulators to control the polymer assembly-disassembly dynamics between monomeric and filamentous actin (F-actin). As an abundant cytoskeleton protein with numerous protein-protein interactions, actin determines cell shape and functions in a variety of fundamental biological processes such as cell division and cell motility (Dominguez & Holmes, 2011; Pollard, 2016; Pollard & Cooper, 2009).

In defending against infection, actin has direct roles in the organization of epithelial barriers by establishing cell polarity and driving the formation of cell junctions (Cavey & Lecuit, 2009; Raman et al., 2018). It may also interfere with pathogen dissemination by enlisting septin cages to restrict bacterial motility within the cytosol of non-immune cells (Krokowski et al., 2018; Mostowy et al., 2010). Furthermore, actin-based mechanical forces are critical for professional immune cells (*e.g.*, macrophages, neutrophils) to migrate to infected tissues (Fritz-Laylin et al., 2017; Wiesner et al., 2014), for phagocytosis, and antigen presentation (Davidson & Wood, 2020; Rougerie et al., 2013). Although ample evidence supports the coupling of actin regulation to host immunity, understanding how specific actin-binding proteins operate within defined infection contexts may provide insights into unidentified mechanisms of immunity and stimulate the development of therapeutic interventions against microbial pathogens.

During infection, actin is commonly exploited by a broad range of microbes. Extracellular bacteria such as pathogenic *E. coli* (enteropathogenic, enterohemorrhagic) and *Helicobacter pylori* employ secretion systems to translocate effector proteins that modify the actin cytoskeleton of epithelial cells to promote microbial adhesion and disrupt barrier integrity during gastrointestinal colonization (Croxen & Finlay, 2010; Rottner et al., 2005; Selbach et al., 2003; Wessler et al., 2011). Intracellular pathogens such as *Burkholderia spp., Listeria monocytogenes, Rickettsia spp., Shigella spp.,* and the vaccinia virus manipulate host actin to invade cells and disseminate using processes such as actin-based motility (Lamason & Welch, 2017; Pizarro-Cerdá & Cossart, 2006; Stevens et al., 2005; Welch & Way, 2013). The pervasiveness of microbial mechanisms that exist to subvert host actin underscores the importance of actin for pathogenesis. Therefore, determining how specific actin-binding proteins impact microbial virulence is imperative to understand infectious diseases.

Given the complexity of the actin cytoskeleton and diversity of actin-binding proteins, there is continuous interest in characterizing how these proteins can alter the course of infection as they are mobilized during cell-autonomous immunity. One of the most powerful signals triggering cell-autonomous immunity is interferon-gamma (IFN-γ), a type II cytokine expressed during infection and inflammation in vertebrates. It stimulates a broad range of host cells to express thousands of interferon-stimulated genes (ISGs) important for defense against bacteria, parasites, fungi, and viruses (MacMicking, 2012; Schneider et al., 2014). Using genome-wide transcriptomic and computational approaches, we identified dozens of actin-associated ISGs expressed by human immune and stromal cells (*i.e.,* macrophages, fibroblasts) and uncovered novel functional roles for one of the most highly induced ISGs - <u>xi</u>n repeat actin-binding <u>p</u>rotein <u>1</u> (*XIRP1*). XIRP1 was found to be associated with macrophage podosomes and co-opted by cytosolic *Listeria* to exit host cells, leading to enhanced pathogenicity in a murine model of infection described herein.

## Results

### Immune activation by IFN-γ induces numerous human actin-binding proteins

To identify actin-associated genes responsive to immune stimuli and potentially relevant to human microbial infections, we combined whole cell RNA-sequencing (RNA-Seq) with *in silico* data-mining approaches. Fibroblasts were initially chosen as a non-immune cell source given their broad tissue distribution and robust response to IFN-γ (Cheng et al., 1985; Tretina et al., 2019). Here primary human derived Hs27 fibroblasts cells were selected for investigation based on their ability to form flattened polygonal sheets with multiple intercellular contacts for examining pathogen dissemination and their non-cancerous phenotype. Additionally, THP-1 cells were chosen as an immune phagocytic model as they are tractable to CRISPR-Cas9 engineering and a widely used surrogate for human primary macrophages. Hs27 fibroblasts were treated with IFN-γ before isolating RNA, RNA-Seq and data analysis. In parallel, we employed the ARCHS^4^ RNA-Seq data-mining web resource (Lachmann et al., 2018) to identify publicly available RNA-Seq datasets of classically activated THP-1 macrophages (IFN-γ ± lipopolysaccharide).

RNA-Seq analysis of Hs27 fibroblasts revealed a robust transcriptional response 24 h after IFN-γ treatment. We detected a total of 982 differentially expressed genes (DEGs) (fold-change ≥ 2, 5% false discovery rate-adjusted *p* value < 0.05), from which 337 (∼34.3%) were downregulated and 645 (∼65.7%) upregulated (*Figure 1a*). Among the most highly induced ISGs were those encoding cell-autonomous immunity proteins of the guanylate-binding family (*GBP1-6*), proteins involved in antigen presentation (*HLA-DOA, HLA-DRA, HLA-DRB1, 5*), sensors of exogenous-RNA from the oligoadenylate synthase family (*OAS1-3, OASL*), chemokines (*CXCL9-11, CX3CL1*), and other genes with known immune functions (*e.g.*, *IDO1, IRF8, APOL1-4, 6*) (Gaudet et al., 2021; MacMicking, 2012; Tretina et al., 2019). Based on Gene Ontology (GO) classification, we identified a subset of 50 (∼5.1%) fibroblast DEGs associated with actin-related functions (*Figure 1a*). These included genes encoding the highly induced anti-viral protein tetherin (*BST2*), the monocyte/macrophage chemoattractant CX3CL1, and the actin-binding protein XIRP1.

**Figure 1.**
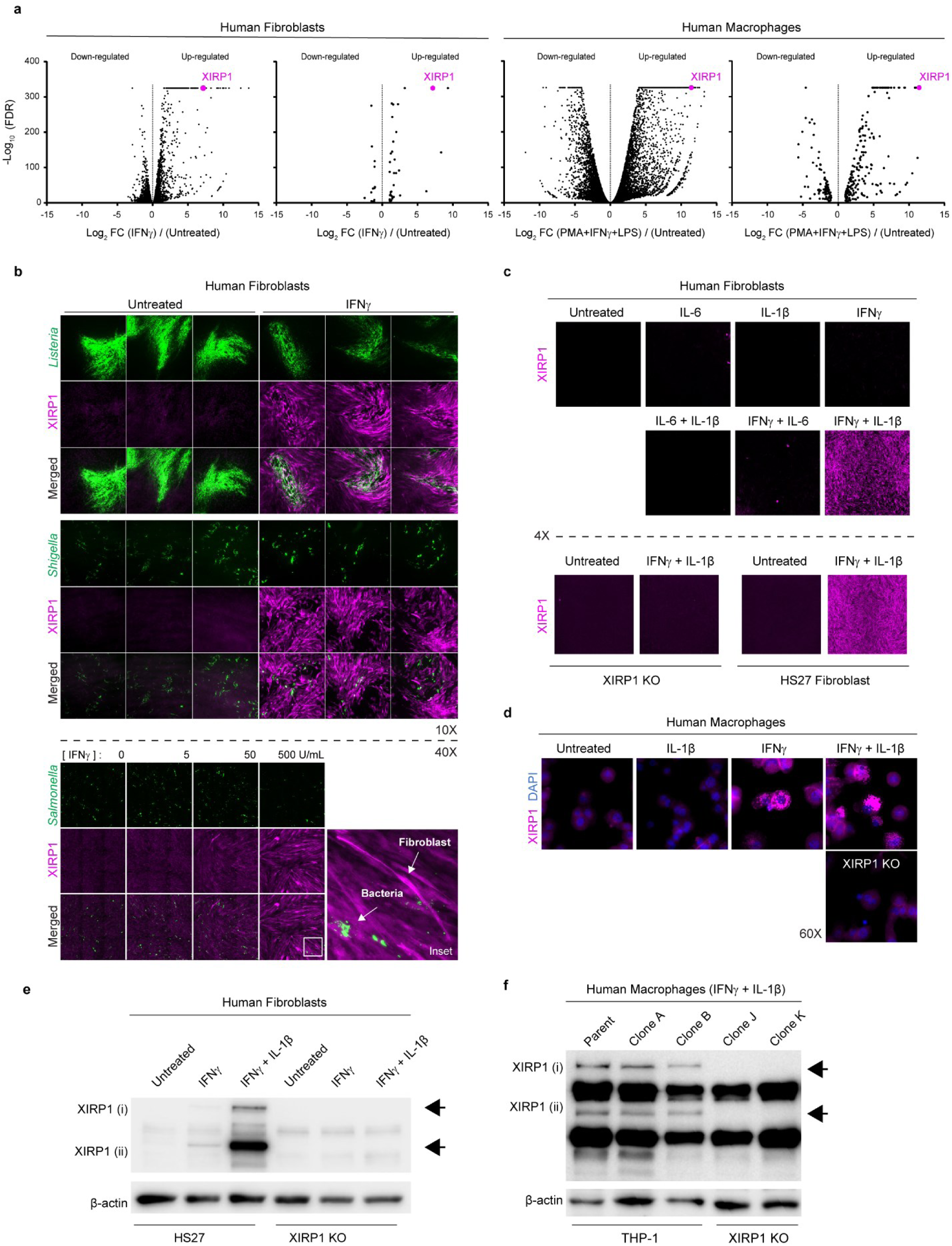
XIRP1 expression by fibroblasts and macrophages. (a) RNA-Seq analysis of Hs27 fibroblasts treated for 24 h with IFN-γ (1000 U/mL) compared to untreated cells and re-analysis of classically activated THP-1 macrophages (PMA differentiated, LPS and IFN-γ treated) compared to untreated THP-1 monocytes (re-analyzed GEO datasets: GSM3729256-9, GSM3729250-2; PMID: 32449925). *XIRP1* is shown in magenta among overall DEGs (left panels) and the subset of actin-associated DEGs (right panels). FDR, 5% false discovery rate-adjusted *p* value; FC, fold change. (b) Immunostaining of Hs27 fibroblast monolayers infected with GFP-expressing *Listeria* (top) or *Shigella* (middle) in the presence or absence of 1000 U/ml IFN-γ. Normal human intestinal myofibroblasts monolayers infected with GFP-expressing *Salmonella* (bottom) at indicated IFN-γ concentrations. (c) Immunostaining of Hs27 fibroblast monolayers and (d) THP-1 derived macrophages 24 h post-treatment with the indicated cytokines. (e) XIRP1 Immunoblots from Hs27 fibroblasts and (f) THP-1 derived macrophages 24 h post-treatment with indicated cytokine.

XIRP1 was ideal for further characterization as it has unknown roles in host immunity, has direct actin-binding activity and is markedly induced by IFN-γ (∼143.3-fold). To gain insight into XIRP1 expression within specialized immune cells, we used the ARCHS^4^ gene search webtool to identify macrophages as a cell type that expresses high levels of XIRP1 under given conditions. ARCHS^4^ allowed the comparison of XIRP1 gene-level counts from a total of 1,933 macrophage RNA-Seq datasets publicly available through the Gene Expression Omnibus (GEO) database (*Figure supplement 1a*). Human monocyte-derived macrophages (HMDM) infected with either *Listeria* or *Salmonella* bacteria were among the samples with the highest normalized counts. Classic M1 macrophage polarization with IFN-γ and lipopolysaccharide (LPS) showed similarly high levels of XIRP1 (*Figure supplement 1a*). Indeed, re-analysis of published datasets via our RNA-Seq pipeline showed XIRP1 as the most induced (∼2745.8-fold) actin-binding protein when THP-1 monocyte cells are compared to M1 polarized THP-1 macrophages (GEO datasets: GSM3729250-2 vs GSM3729256-58; PMID 32449925) (*Figure 1a*).

Despite differences in cell type, Hs27 fibroblasts and THP-1 macrophages shared other common transcriptional responses to IFN-γ treatment. ISGs highly induced in both cell types include *GBP1-6*, *OAS1-3*, *OASL*, *IDO1*, *APOL1-4*, *APOL6*, and *HLA*-genes (*Supplementary Table 1 and 2*). Besides XIRP1, highly induced genes encoding actin-associated proteins include *BST2, ICAM1,* and *MYPN* (*Supplementary Table 1 and 2*). Macrophage actin-associated DEGs amounted to 303, exceeding the 50 DEGs of fibroblasts. However, the proportion was similar with respect to total DEGs in each cell type (∼3.7% vs ∼5.1%). The greater number of overall DEGs in macrophages likely reflects the specialized role of these cells in immunity, but also a more complex immune stimulation as THP-1 cells were first differentiated to macrophages and then polarized with IFN-γ and LPS.

### Human *XIRP1*: An IFN-γ-induced gene enhanced by IL-1β or bacterial signals

In vertebrates ranging from sea lamprey to humans, expression of XIRP1 homologs is associated with homeostatic and pathological heart function (Grosskurth et al., 2008; Wang et al., 2014). Additional roles in development and skeletal muscle repair have also been noted in zebrafish and other organisms (Braun et al., 2019; Otten et al., 2012). We were therefore surprised to observe such profound XIRP1 induction in response to IFN-γ and other classical immune stimuli (*Figure 1a-f*). Phylogenetic and gene synteny analysis lent further support for a potential role of XIRP1 in host immunity. Searching within genetic regions flanking *XIRP1* identified linkages with MYD88 across numerous vertebrates (*Figure supplement 1b*); this innate immune adapter is critical for toll-like receptor (TLRs) and IL-1β signaling (Deguine & Barton, 2014). Within mammals, *XIRP1* was also surrounded by chemokine receptor genes important for macrophage chemotaxis (*i.e., CX3CR1*, *CCR8*). Indeed, in humans both the *XIRP1* and *MYD88* loci reside within two chemokine receptor clusters on chromosome 3. The dramatic reduction in xin repeat domains noted for human and other mammalian XIRP1 also suggested new roles beyond host musculoskeletal functions (Grosskurth et al., 2008).

Directly testing immune responsiveness of XIRP1 further hinted at a role in host defense. We performed immunoblots and immunostaining of fibroblast and macrophage samples in response to different immune and infectious stimuli. Here we generated CRISPR-Cas9 stable XIRP1 deletions (XIRP1 KO) to confirm antibody specificity and for use in functional studies. Pre-treatment of fibroblast monolayers with IFN-γ followed by infection with diverse intracellular bacteria (*i.e., Listeria*, *Shigella*, and *Salmonella*) led to robust XIRP1 expression (*Figure 1b*). In the absence of infection, however, IFN-γ, IL-1β, or IL-6 alone did not yield detectable XIRP1 protein, despite mRNA transcripts in our RNA-Seq analysis. This suggests additional infection-induced signals were needed for full expression. Indeed, combined treatment with IFN-γ and IL-1β led to robust XIRP protein levels, suggesting both STAT1 and nuclear factor kappa B (NF-κB) signals converge on the human *XIRP1* promoter (*Figure supplement 1c*). Notably, in phorbol myristate acetate (PMA)-derived THP-1 macrophages IFN-γ was sufficient for XIRP1 expression (*Figure 1d*). Here PMA activation of NF-κB probably produces sufficient autocrine IL-1β to trigger XIRP1 expression in combination with IFN-γ (*Supplementary Table 2*) (Hellweg et al., 2006; Takashiba et al., 1999). Collectively, these data show stimulation of multiple pathways is required for optimal XIRP1 expression in both immune and non-immune cells.

### IFN-γ-induced XIRP1 localizes to macrophage podosomes

To identify potential roles for XIRP1 in host immunity we first focused on characterizing its cellular localization within IFN-γ-activated macrophages. Fluorescent confocal microscopy revealed endogenous XIRP1 organized in an unusual, bespeckled pattern at the base of cells near the underlying substrate (*Figure 2b-c*). These clusters resembled actin-rich adhesion structures in macrophages known as podosomes (Wiesner et al., 2014). Indeed, staining for F-actin found XIRP1 forms a unique toroidal pattern around a central F-actin core (*Figure 2b, c, e*). The prototypic conical shape of podosomes was revealed in orthogonal views and 3D reconstructions of images collected via confocal microscopy (*Figure 2c, d*).

**Figure 2.**
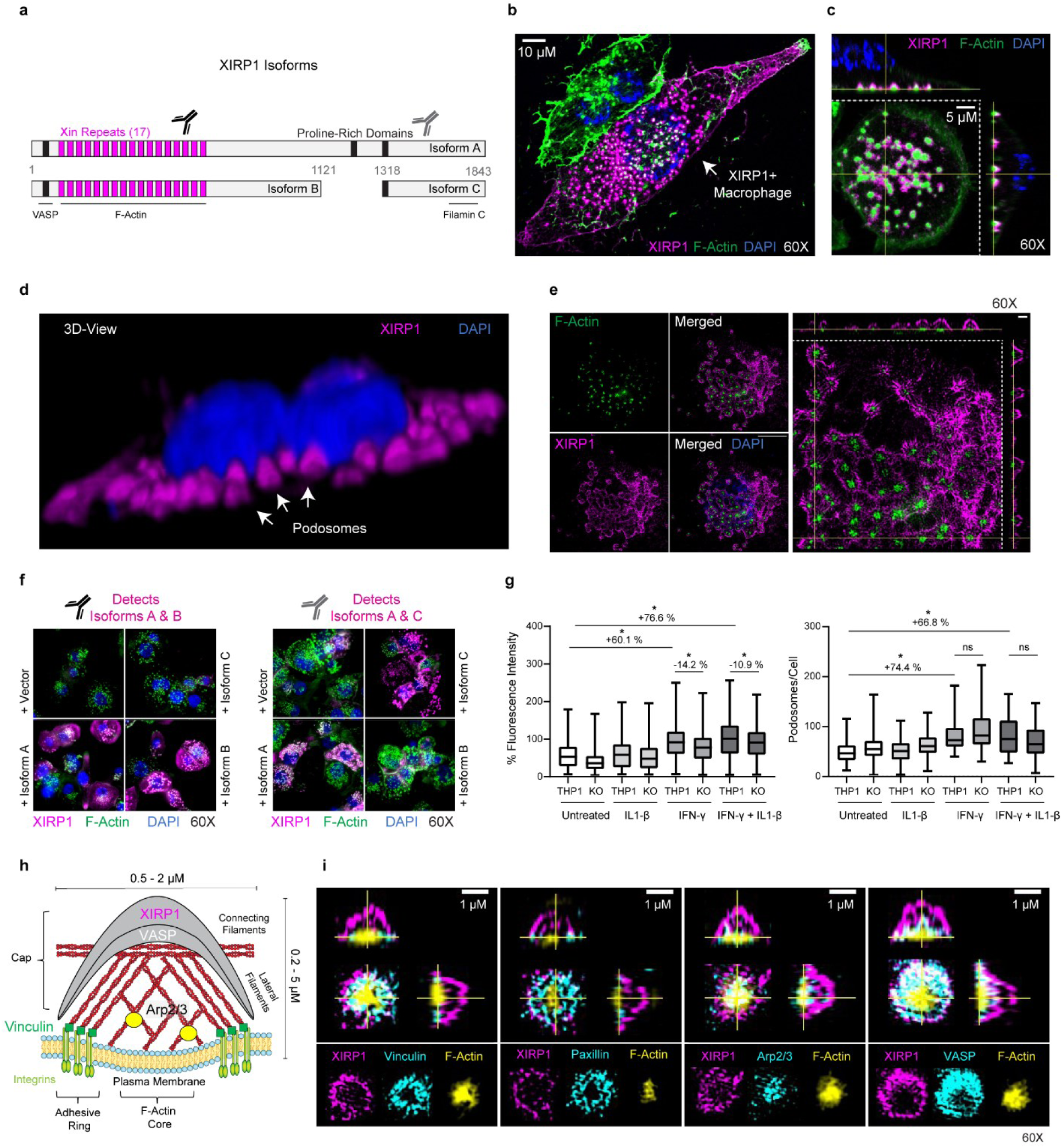
XIRP1 localization in macrophages podosome networks and single podosomes. (a) Diagram of the three human XIRP1 isoforms shows the actin-binding xin repeat domains, proline-rich domains and selected XIRP1 regions previously shown to interact with VASP, F-actin, and Filamin C. The epitopes detected by the anti-XIRP1 antibodies are indicated. (b) A representative single z-slice confocal microscopy image of immunostained THP-1 macrophages shows XIRP1 localization to sub-cellular podosomes (magenta and green specks). (c) orthogonal views and (d) 3-D rendering of confocal microscopy images show dozens of conical XIRP1 structures distributed at the base of macrophages. (e) Super-resolution structured-illumination microscopy of a single cell immunostained for XIRP1(left) and orthogonal views of the macrophage podosome network (right). (f) Epi-fluorescent microscopy images of XIRP1-knockout macrophages and derived cells expressing individual isoforms upon retroviral transduction. Immunostaining with isoform-specific antibodies shows podosome localization of individual XIRP1 isoforms. (g) Measurement of F-actin fluorescence intensity in podosomes (n ≥ 1884) and (e) number of podosomes per cell (n ≥ 30 cells) in THP-1 macrophages and XIRP1-knockout cells. \**p* < 0.05, unpaired t test. (h-i) Super-resolution structured-illumination microscopy of single podosomes immunostained for XIRP1 and indicated podosome markers. Orthogonal views are shown (top) along with individual channels from a single z-slice through the base of the podosome (bottom). The model shows the localization of XIRP1 extending from the podosome cap to the adhesion ring along the lateral actin filaments. Dimensions on the model are based on approximations previously reported.

Since the XIRP1 protein is expressed as three alternatively spliced isoforms of varying length and domain composition (*Figure 2a*), we investigated which isoform(s) assembled in podosomes. Isoforms A (a.a. 1-1843), B (a.a. 1-1121), and C (a.a. 1318-1843) were individually re-introduced into XIRP1 KO human macrophages via retroviral transduction. Transductions with shorter isoforms B and C were more efficient than those with the full-length isoform A which showed a mixed XIRP1-positive and -negative macrophage population (*Figure 2f, lower-left panel*s). Nonetheless, all three isoforms were recruited to podosomes as shown by immunostaining with isoform-specific antibodies (*Figure 2f*). Isoforms A and B contain xin repeat domains which directly bind F-actin as well as a proline-rich domain that physically interacts with the actin-binding protein VASP; isoform C does not contain xin repeats but does include domains that engage Filamin C and other actin-binding proteins (van der Ven et al., 2006). Since podosomes are complex adhesion structures composed of F-actin and numerous associated proteins, it is likely that XIRP1 isoforms are recruited through both actin-binding xin repeats and other domains that engage podosome accessory proteins.

Super-resolution structured illumination microscopy (SIM) verified XIRP1 encapsulation of the F-actin core at the base of individual podosomes and throughout the macrophage podosome network (*Figure 2e*). XIRP1 forms a dome-shaped cap that extends down into the adhesion ring composed of vinculin and paxillin (*Figure 2h, i*). Because XIRP1 is excluded from the branched F-actin core containing Arp2/3 and instead is closely associated with the bundling protein, VASP, the precise location of XIRP1 seems to extend along the lateral, linear actin filaments of podosomes.

Examination of XIRP1 KO macrophages found a significant reduction in F-actin content in podosomes specifically after IFN-γ treatment (*Figure 2g*); loss of XIRP1 did not, however, prevent podosomes being formed (*Figure 2f, upper-left panel*). Hence XIRP1 appears to contribute to alterations in existing podosome architecture specifically upon immune activation, consistent with its cytokine-induced expression profile.

### Cytosolic *Listeria* co-opts XIRP1 for actin-based motility in macrophages

Having found XIRP1 is a component of the actin-based podosome network in IFN-γ activated macrophages, we wondered whether pathogens reliant on the actin-based network may be subject to XIRP1-mediated control in immune-activated cells or instead use these structures as platforms for locomotive escape within the cytosol. *Listeria monocytogenes* is an intracellular pathogen that evades lysosomal degradation by escaping into the cytoplasm where it hijacks cell actin for motility and cell-to-cell dissemination (*Figure 3a*) (Radoshevich & Cossart, 2018). The ActA virulence factor of *L. monocytogenes* directly recruits the ARP2/3 complex and VASP to polymerize host actin and propel bacteria (Lambrechts et al., 2008). Since XIRP1 is a known interacting partner of VASP, it is possible XIRP1 may also interact with *L. monocytogenes* to retard or facilitate its spread.

**Figure 3.**
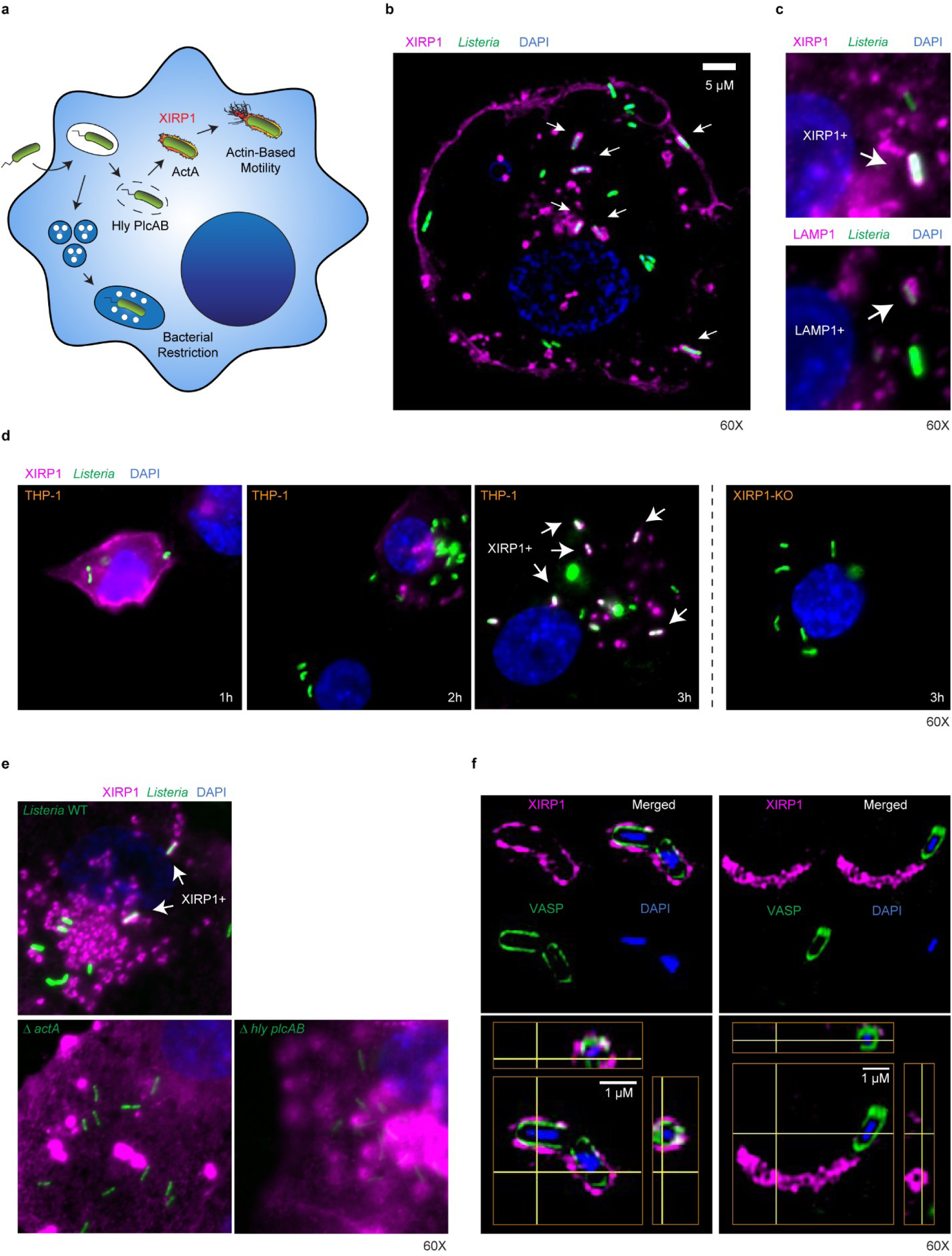
XIRP1 recruitment to the surface of intracellular *Listeria* in THP-1 macrophages. (a) Model shows *Listeria* invasion of the macrophage cytoplasm and actin-based motility. (b) Confocal microscopy of infected THP-1 macrophages (single z-slice, 3 h post-infection). (c) Immunostaining against lysosomal LAMP1 and XIRP1. (d) Immunostaining of infected THP-1 macrophages fixed at indicated time-points post-infection. (e) Immunostaining of THP-1 macrophages infected with wildtype *Listeria* or mutants deficient in actin-based motility (Δ*actA*) or phagosome rupture (Δ*hly plcAB*). (f) Super-resolution structured-illumination microscopy of replicating (left panels) and motile (right panels) *Listeria* in THP-1 macrophages (single channels, top; orthogonal views, bottom). **Figure 3—video 1.** **XIRP1 localizes to the surface of replicating *Listeria* in THP-1 macrophages** Live-cell widefield microscopy of THP-1 macrophages expressing XIRP1B-mCherry ∼2.5 h post-infection with GFP-expressing *Listeria.* Bar: 10 μm **Figure 3—video 2.** ***Listeria* escapes the XIRP1 coat after replication in THP-1 macrophages** Live-cell widefield microscopy of THP-1 macrophages expressing XIRP1B-mCherry ∼2.5 h post-infection with GFP-expressing *Listeria.* Bar: 10 μm **Figure 3—video 3.** **XIRP1 localizes to the actin-tails of motile *Listeria* in THP-1 macrophages** Live-cell widefield microscopy of THP-1 macrophages expressing XIRP1B-mCherry ∼2.5 h post-infection with GFP-expressing *Listeria.* Bar: 10 μm **Figure 3—video 4.** **XIRP1 localizes near the plasma membrane as *Listeria* escapes THP-1 macrophage cells** Live-cell widefield microscopy of THP-1 macrophages expressing XIRP1B-mCherry ∼2.5 h post-infection with GFP-expressing *Listeria.* Bar: 10 μm

To first determine whether XIRP1 is recruited to intracellular *Listeria*, we performed immunostaining of endogenous XIRP1 during infection of IFN-γ-activated macrophages. At early time points post-infection (1-2 h p.i.), XIRP1 was absent from *L. monocytogenes* expressing Green Fluorescent Protein. However, by 3 h p.i. it was readily detected around *Listeria* in both epifluorescence and confocal microscopy (*Figure 3b, d*). Importantly, the immunofluorescent signal was missing in XIRP1 KO macrophages, underscoring antibody specificity and not fluorescent bleed-through. Moreover, XIRP1-positive bacteria did not co-localize with the lysosomal marker LAMP1, while LAMP1-positive bacteria in turn did not co-localize with XIRP1 (*Figure 3c*). Hence XIRP1 is specifically recruited to *L. monocytogenes* that have escaped their entry vacuole to replicate in the cytosol. This was validated using bacterial mutants lacking the virulence factors listeriolysin plus phospholipase A and B (Δ*hly*Δ*plcAB*) that are obligate for vacuolar rupture; these triple mutants did not recruit XIRP1 (*Figure 3e*). Similarly, Δ*actA* mutants which access the cell cytoplasm but cannot recruit host actin also failed to colocalize with XIRP1 (*Figure 3e*). Collectively, the results indicate that *Listeria* recruits XIRP1 upon accessing the cytoplasm in an ActA-dependent manner.

Super-resolution SIM and live widefield microscopy further delineated XIRP1 association with cytoplasmic *Listeria* in THP-1 macrophages. Notably, XIRP1 specifically colocalized with replicating and individual motile bacterium (*Figure 3f and Figure 3—video 1-4*). XIRP1 formed an outer perimeter surrounding VASP on the surface of *Listeria* (*Figure 3f, left*) and a characteristic bend at the site of septation was observed during bacterial replication. Bacterial escape then proceeded through the midpoint of the XIRP1 encasing, suggesting a possible break in the actin coatomer (*Figure 3—video 2*). Upon initiation of actin-based motility, XIRP1 was redistributed to the actin-tail where it forms a hollow tube as VASP remains associated to the surface of *Listeria* (*Figure 3f, right; Figure 3—video 3*). XIRP1 enrichment was also observed behind *Listeria* as it escaped macrophages (*Figure 3—video 4*). *L. monocytogenes* is a pathogen that hijacks numerous actin-binding proteins to invade cells and disseminate in host tissues. Since XIRP1 is mobilized by host immunity in response to microbes, *Listeria* likely adapted to utilize XIRP1 as a mechanism to escape antimicrobial host cells such as macrophages.

## Impaired *Listeria* dissemination in *Xirp1*^-/-^ mice

During infection, XIRP1 is induced in fibroblasts, macrophages, and other cell types. As a host factor involved in cell adhesion processes such as podosome- or focal adhesion-mediated migration, XIRP1 could benefit the host by facilitating macrophage recruitment to sites of microbial infection. As a hijacked actin-binding protein, XIRP1 may be detrimental to the host by promoting bacterial dissemination. Either outcome was evaluated during *Listeria* infection. Wildtype C57BL/6NJ or XIRP1-deficient (*Xirp1*^-/-^) mice were challenged intraperitoneally and survival monitored for 14 days p.i. Dissemination to secondary organs was assayed via colony forming units (CFU) in the liver, spleen, heart, and brain at 4 days p.i. *Xirp1*^-/-^ mice were significantly more resistant than wildtype mice and had median bacterial burdens that trended ∼1 Log_10_ lower in most organs (*Figure 4*). The heightened resistance and reduced dissemination in *Xirp1*^-/-^ mice suggests XIRP1 is subverted by *Listeria* to support actin-based motility and cell-to-cell spread. It remains formally possible that other pathogens which do not use actin-based motility may be controlled via host defense processes in an XIRP1-dependent manner, for example, macrophage migration to infectious foci. However, these require additional testing in future.

**Figure 4.**
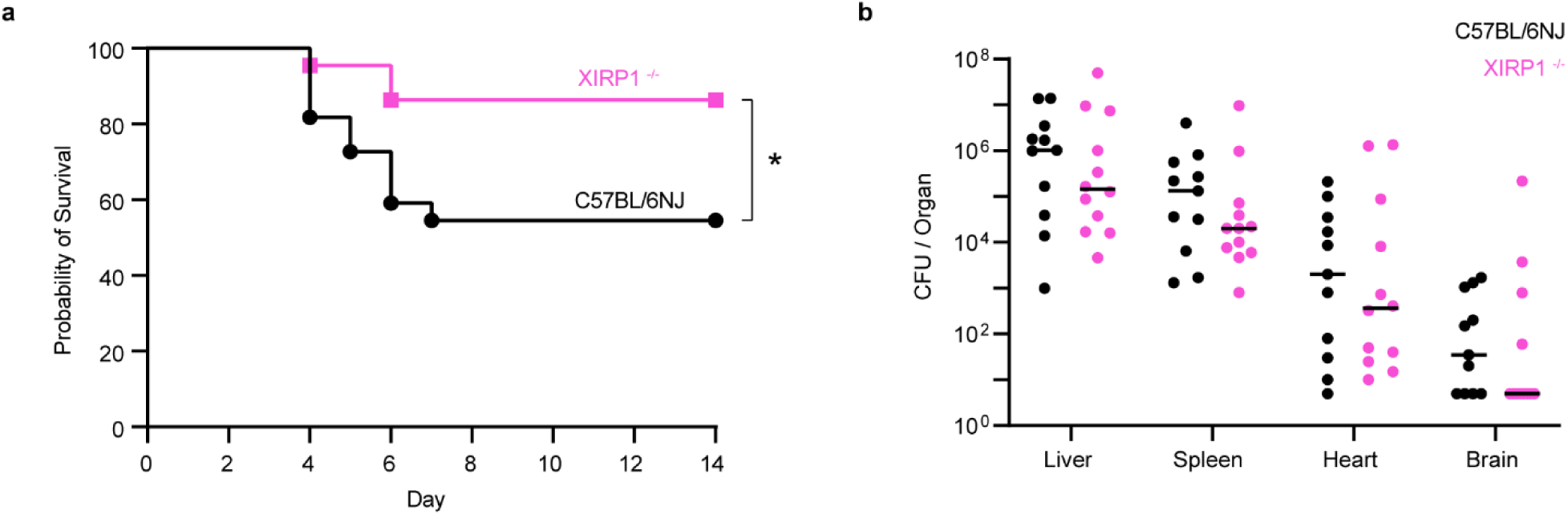
XIRP1 is detrimental to the host during *Listeria* infection. (a) Mouse survival after intraperitoneal infection with *Listeria* (7.5×10^5^ CFU per mouse). \**p* < 0.05, log-rank (Mantel-Cox) test, n = 22 mice per group. (b) Bacterial burden 4-days post-infection per indicated organ. Horizontal line shows median CFU, n ≥ 11 mice per group. From thirteen total mice per group, two C57BL/6NJ and one XIRP1^-/-^ mice did not survive 4-days post-infection and were excluded.

## Discussion

IFN-γ elicits large-scale defense programs central to cell-autonomous immunity in humans and other vertebrates against a wide range of microbial pathogens (MacMicking, 2012). Here we show that actin-binding proteins constitute an important fraction of ISGs mobilized by host immunity and can impact the outcome of infection. Proteins such as XIRP1 may provide mechanical support for endothelial transmigration or tissue infiltration, as podosomes are known to initiate invasive cell protrusions through the vasculature and deliver proteases that degrade the extracellular matrix (Carman et al., 2007; Linder, 2007; Van Goethem et al., 2011). Although we focused on macrophage podosomes, we also observed XIRP1 at focal adhesions (*Figure supplement 2*) and other actin-rich structures in fibroblasts. These stromal cells have underappreciated roles in host-microbe interactions, yet they are known to regulate immune processes and even restrict microbial pathogens (Davidson et al., 2021; Gaudet et al., 2016; Gaudet et al., 2021; McCormack et al., 2013).

Many ISGs protect the host against infection, however, pathogens can adapt to circumvent immunity and even exploit host defenses. Our results suggest XIRP1 is another host protein hijacked by *Listeria*, a master manipulator of host actin. Our observations indicate that XIRP1 is recruited at a specific stage of the *Listeria* intracellular lifecycle once it escapes into the cytosol. Here ActA is critical as it assembles host proteins to drive *Listeria* actin-based motility. These include the Arp2/3 actin nucleation complex, profilin, G-actin, and Ena/VASP proteins involved in filament elongation (Kocks et al., 1992; Lambrechts et al., 2008; Welch et al., 1997). XIRP1 recruitment to *Listeria* was also dependent on ActA but it is unclear whether it occurred directly or indirectly via VASP. The N-terminal proline-rich domain of XIRP1 is likely to be involved given it directly interacts with the EVH1 domain of VASP. We found both proteins closely apposed around *Listeria* and in podosomes. Notably, actin recruitment to *Listeria* and actin-based motility was still evident in *Xirp1* KO macrophages, suggesting other accessory proteins provide a degree of compensation. Future studies should clarify whether XIRP1 is only co-opted by *Listeria* or by other pathogens that manipulate host actin (*e.g.*, *Burkholderia spp., Rickettsia spp., Mycobacteria marinum, Shigella spp, Vaccinia virus)*.

Our work provides new insight into a role of XIRP1 in host-microbe biology, yet it remains unclear how XIRP1 affects the mechanical properties of the actin macromolecular structures to which it binds. One possible function for XIRP1 may include its filament bundling activity which could enhance actin-tail stiffness and allow for greater force generation. This might be particularly important for bacterial dissemination in tissues where cell-matrix and cell-cell adhesions act as mechanical barriers against colonization. Cryo-electron tomography has shown bundles of nearly parallel actin filaments on the surface of *Listeria* and its actin-tails (Jasnin et al., 2013). Consistently, XIRP1 was found to be excluded from the branched actin core in podosomes, suggesting an association with the outer linear filaments. Akin to *Listeria* actin-tails, XIRP1 may also stiffen the upper dome-shaped structure on podosomes to allow for greater protrusive forces as macrophages push against a substrate. More detailed biophysical studies on *Listeria* motility and cell podosomes should clarify the role of XIRP1 in these novel biomechanical functions.

Xin repeat-containing proteins are recently discovered and most studies to date are limited to skeletal muscle and cardiac functions. Our findings expand the repertoire of host cell types known to express XIRP1 and suggest intimate involvement of this new actin-binding protein with the cell adhesion machinery during infection. Importantly, pronounced XIRP1 expression in immune cells strongly implicates activities beyond musculoskeletal homeostasis. *In vivo* challenge experiments clearly reveal such non-canonical activities operate *in vivo*. Future work on XIRP1 conformers could provide a structural basis for therapies against infection, autoimmunity, cardiovascular disease and even cancer.

## Materials and methods

### Bacterial strains and growth conditions

Strains for cell culture experiments included GFP-expressing *Listeria monocytogenes* 10403S (DHL1039), Δ*actA* and Δ*hly* Δ*plcAB* mutants (H. Agaisse). *Listeria* was grown overnight in brain heart infusion (BHI) broth (HiMedia Laboratories, LLC) at 28°C with shaking (225 rpm). For animal experiments, streptomycin-resistant *Listeria monocytogenes* 10403S (J. J. Woodward laboratory) was sub-cultured in fresh BHI media (1:50 dilution) and grown to OD_600_ = 0.6 at 37°C with shaking (250 rpm). *Shigella flexneri* M90T (F. Randow) and *Salmonella* Typhimurium 1344 (J. Galan) expressing GFP from pFPV25.1 (Addgene) were grown in Luria-Bertani (LB) broth (Thermo Fisher Scientific) containing carbenicillin (100 μg/ml) at 37°C overnight with shaking (250 rpm). For fibroblast infections, *Shigella* was sub-cultured in fresh LB (1:100) and grown to OD_600_ = 0.6 at 37°C with shaking (250 rpm). *Salmonella* was sub-cultured to OD_600_ ∼1.0.

### Cell lines and growth conditions

Cell cultures were maintained at 37°C in a 5% CO_2_ incubator. Normal human fibroblast Hs27 cells (ATCC, CRL-1634) were cultured in DMEM (Thermo Fisher Scientific) supplemented with 10% heat-inactivated fetal bovine serum (FBS) (Thermo Fisher Scientific). Primary human intestinal myofibroblast (Lonza Group AG, CC-2902) were maintained in SmGM Medium with growth factors (Lonza Group AG). Human THP-1 monocytes (ATCC, TIB-202) were grown in RPMI 1640 Medium with ATCC modification (Thermo Fisher Scientific) supplemented with 10% FBS and 50 μM 2-mercaptoethanol (Millipore Sigma). Macrophages were generated by differentiating THP-1 monocytes with 100 ng/ml phorbol 12-myristate 13-acetate (PMA) (Millipore Sigma, P8139) for 24 h. Macrophages in podosome experiments were then activated with IFN-γ for an additional 24 h. For bacterial infections, PMA was removed for 24 h prior to cytokine treatment.

Unless otherwise indicated, cells were treated for 24 h with either 50 ng/ml IFN-γ (R & D Systems), 10 ng/ml IL-1β (R & D Systems),100 ng/ml IL-6 (R & D Systems), or a combination of cytokines.

### Construction of plasmids

For CRISPR-knockout plasmids, two oligonucleotide pairs carrying sgRNA sequences were designed to target XIRP1 and ordered (Integrated DNA Technologies) for cloning into pSpCas9(BB)-2A-Puro (pX459) V2.0 (Addgene, 62988). A pair was designed targeting a region upstream of the translational start site and consisted of 5’-CACC<u>GGG AGTGACCATGGTACCAC</u>-3’ and 5’-AAAC<u>GTGGTACCATGGTCACTCCC</u>-3’. The second pair was designed to target downstream of the translational stop site and consisted of 5’-CACC<u>GATTCACACATGTCGATGCGT</u>-3’ and 5’-AAAC<u>ACGCATCGACATGTGTGAATC</u>-3’. Oligonucleotide pairs (10 μM) were hybridized in T4 Ligase buffer using a thermocycler where samples were incubated at 95°C for 5 min, 95°C for 15 sec, and 92.5°C for 15sec. The last two incubations were repeated for 28 cycles, lowering the temperature by 2.5°C every cycle. Plasmid pX459 was digested with *Bbs*I (New England Biolabs) and purified with the Wizard SV Gel and PCR Clean-Up System (Promega) after DNA electrophoresis. Ligation of cut pX459 (100 ng) and diluted hybridized oligonucleotides (1:200) was performed with T4 Ligase in 20 μl reactions by incubating for 30 min at room temperature. Plasmid constructs were electroporated into *E. coli* DH10B ElectroMAX (Thermo Fisher Scientific) bacteria and verified by Sanger sequencing (Genewiz, Inc).

Retroviral vectors introducing individual XIRP1 isoforms were generated with pMSCVpuro (Clontech Laboratories, 631461). Since the XIRP1 coding region is entirely within exon 2, PCR amplification was directly done from gDNA. Isoform A was amplified with primer 5’-gaattagatctctcgaggttCCACCATGGCCGACACCCAGACACAGGTG-3’ and 5’-cctacccggtaga attcgttTCACTGGGCAGCTGGCTGGGAGTAGCTGCA-3’; isoform B with 5’-gaattagatctctcgag gttCCACCATGGCCGACACCCAGACACAGGTG-3’ and 5’-ctacccggtagaattcgttTCACAGCAGCTTTCTGGGGGCTGCTGGGATCCGGGGATCAC-3’; isoform C with 5’-gaattagatctctcgagg ttCCACCATGCCCCCAAAGAAGAAGCCGCAGCTG-3’ and 5’-cctacccggtagaattcgttTCACTGGGCAGCTGGCTGGGAGTAGCTGCA-3’. Fragments were cloned into *Hpa*I-digested pMSCVpuro using the Gibson Assembly Cloning Kit (New England Biolabs, E5510S). Plasmids were transformed into *E. coli* NEB 5-alpha and the purified plasmids verified by Sanger sequencing (Genewiz, Inc).

To construct the XIRP1-mCherry translational reporter, the retroviral vector pMSCV-IRES-mCherry FP (addgene, 52114) was modified by fusing the sequence from XIRP1 (isoform b) to the N-terminus of mCherry. The XIRP1 (isoform b) fragment was PCR amplified with a C-terminal linker (GlyGlyGlyGlySer) added with primers 5’-CACC<u>ATG</u>GCCGACACCCAGACA CAG-3’ and 5’-<u>GGAGCCCCCTCCGCC</u>CTTTCTGGGGGCTGCTGGGATC-3’. To allow for Gibson cloning, a second PCR was performed adding complementary regions to the vector with primers 5’-tttgaaaaacacgataatacCACCATGGCCGACACCCAG-3’ and 5’-tcctcctcgcccttgctcacG GAGCCCCCTCCGCCCTT-3’. The resulting fragment was then incorporated to *Nco*I-digested pMSCV-IRES-mCherry using the Gibson Assembly Cloning Kit (New England Biolabs, E5510S). The construct was transformed into *E. coli* NEB 5-alpha and the purified plasmid verified by Sanger sequencing (Genewiz, Inc).

### CRISPR-Cas9 mutagenesis

Gene-knockout in Hs27 fibroblasts was performed by transfecting pX459 derived plasmids with WI-38 Cell Avalanche Transfection Reagent (EZ Biosystems, LLC). A day prior to transfections, cells were seeded in DMEM (10% FBS) at 2 x 10^5^ cells per well in 6-well plates. The two plasmids delivering sgRNAs targeting XIRP1 were mixed 1:1 and 4 mg of total endotoxin-free DNA was added to 200 μl OptiMEM containing 6.4 μl of transfection reagent. The mixture was briefly vortexed, incubated for 15 min at room temperature, and 150 μl was added per well. Plates were centrifuged at 300 x g for 5 min, then incubated for 5 h at 37°C in a 5% CO_2_ incubator. The media was replaced to remove transfection reagent, and puromyocin (1-2 μg/ml) selection was started 24 h later. Selection was held for 24-48 h and media was then replaced with DMEM (10% FBS) without puromycin to allow cells to recover for 24-48 h. Adherent cells were dislodged with StableCell Trypsin Solution (Millipore Sigma) and diluted (∼3 cells per 200 μl) in fresh DMEM (20% FBS) containing 33% spent media from Hs27 cultures (80-95% confluent). Clone isolation was performed by limited dilution in 96-well plates using 200 μl of cell suspension per well. The media was replaced every fifth day for ∼25 days or until clones were clearly visible using 4 x magnification. To screen clones, DNA extract was prepared from an aliquot of cells with QuickExtract DNA Extraction Solution (Lucigen Corporation). Only wells containing single clones were screened using PCR primers flanking the Cas9 cut sites and by Sanger sequencing (Genewiz, Inc). Cut site 1 was screened using PCR primer pair 5’-CTAGCTCAGACATTCTCAG TC-3’ and 5’-GGTAGTAAACAGGGCTCAG-3’; the second cut site was screened with 5’-CAGGACTGAAGTGGGTAC-3’ and 5’-GGAACACATACTGGAACAGATG-3’.

As THP-1 monocytes are difficult to transfect, gene-knockout was performed by electroporation of TrueCut Cas9 Protein V2 (Thermo Fisher Scientific) complexed to TrueGuide Synthetic guide RNAs (Thermo Fisher Scientific). Gene-disruption was performed with two sgRNAs, one targeting upstream of the translational start site and another within Exon 2. Sequences of the 1-piece chemically modified synthetic gRNAs were 5’-GGGAGUGACCAUGGUAC-3’ and 5’-GAGACGUUCGUGCAGCCCGC-3’. A total of 20 μg Cas9 was mixed with 120 pmoles of each sgRNA in a 6 μl volume and incubated for 10 min at room temperature. Complexed Cas9-sgRNA was added 49 μl of BTXpress electroporation solution (BTX Molecular Delivery Systems) and incubated for 10 min at room temperature. Media was removed from THP-1 monocytes cultures (5-6 x 10^5^ cells/ml) and cells were washed once with DPBS (Thermo Fisher Scientifics). A total of 2 x 10^6^ cells were resuspended in 50 μl of BTXpress solution and the Cas9-sgRNA complex was added. A 100 μl aliquot of the mixture was transferred to a 0.2 cm gap cuvette and electroporation was performed at room temperature using a square wave protocol (140 V, 1 pulse of 10 msec; field strength 750V/cm) with an ECM 830 Square Wave Electroporation System (BTX Molecular Delivery Systems). Mouse T cell Nucleofector media (100 μl) was added and cells were immediately transferred to a 24-well culture dish containing 1 mL of additional media. Cells were cultured for 72 hours prior to plating for clone isolation. Limited dilution was performed as with Hs27 cells, but 96-well round bottom plates were used instead of flat bottom, and cells were diluted in RPMI media containing 33% spend media from THP-1 suspension cultures (density 6-8 x 10^5^ cells/ml). Clones were PCR-screened for deletions with primers 5’-CTAGCTCAGACATTCTCAGTC-3’ and 5’-GTCTCAAA GATCCAGCGAG-3’ and verified by Sanger sequencing (Genewiz, Inc) with 5’-CTTCACTGTC TTTTTCACATCAC-3’ and PCR primers.

### Fibroblast and macrophage infections

Hs27 fibroblast cells were seeded in 4-well chambered cover glass (Cellvis, C4-1.5H-N) at 0.6 x 10^5^ cells per chamber in 0.5 ml DMEM. Media was replaced with DMEM containing 50 ng/ml IFN-γ or without cytokine and cells were incubated for 24 h. *Listeria* were grown as previously indicated, 1 ml of 18-24 h overnight culture was centrifuged at 10,000 x g for 3.5 min, washed once with phosphate buffer saline (PBS) (Thermo Fisher Scientific), then resuspended at ∼2.5 x 10^9^ CFU/ml in pre-warmed DMEM. Serial dilutions (1:1600, 1:3200, 1:6400) were prepared and fibroblast monolayers were infected with 0.5 ml of diluted bacteria (MOI ∼ 13, 6.5, 3.3). Chambered cover glass was centrifuged at 1000 x g for 5 min prior to incubating for 1 h at 37°C in a 5% CO_2_ incubator. To remove extracellular bacteria, chambers were washed twice with DMEM and media containing 10 μg/mL gentamycin was added. Bacterial infection foci were allowed to develop for 24 h prior to fixation with 4% paraformaldehyde (PFA) solution (Santa Cruz Biotechnology, Inc) for 20 min. *Shigella* infections (MOI ∼ 5.7, 1.1, 0.2) were performed similarly with serial dilutions prepared from OD_600_ = 0.6 cultures. Primary intestinal myofibroblast treated with indicated IFN-γ concentrations were infected with *Salmonella* (MOI ∼20) prepared from OD_600_ ∼1.0 cultures.

### Immunoblots

Fibroblasts were seeded (∼2.5 x 10^5^ cells/well) in 6-well plates containing 2 ml DMEM and allowed to adhere for 24 h or until ∼80-90% confluent. Media was replaced with DMEM containing indicated cytokines and cells incubated for 24 h. For THP-1 cells, monocytes (6 x 10^5^ cells/well) were differentiated to macrophages in 6-well plates for 24 h in 2 ml RPMI containing PMA. Media was replaced with RPMI without PMA and macrophages were incubated for an additional 24 h. Macrophages were then treated with indicated cytokines for 24 h. Cells in each well were lysed in ∼120 μl sample buffer (50mM Tris HCl pH 6.8, 0.02% bromophenol blue, 12.5 mM EDTA, 2% SDS, 10% glycerol, 4% 2-mercaptoethanol) and boiled for 5-15 minutes.

A 20 μl volume of sample was loaded per well of polyacrylamide gel (4% stacking, 8% resolving) and proteins separated by SDS-PAGE (200 V, 50 min). Gels were blotted onto PVDF membranes overnight at 4°C in Towbin buffer (30 V, ∼16 h). PVDF membranes were blocked for 1 h at room temperature in blocking buffer (PBS, 5% non-fat milk, 0.1% tween-20). Primary rabbit polyclonal anti-XIRP1 antibody (Thermo Fisher Scientific, PA5-53487) was added at a 1:5,000 dilution or mouse anti-β-actin [AC-15] (abcam, ab6276) at a 1:10,000 dilution and incubated at room temperature for 2 h with rocking. PVDF membranes were washed for 10 min three times in PBS-T (PBS, 0.1% tween-20). Secondary HRP-conjugated anti-mouse (GE Healthcare, NA931) or anti-rabbit (GE Healthcare, NA934) were added at 1:10,000 dilution for 1.5 h. PVDF membranes were washed for 10 min three times with PBS-T. To image blots, Clarity Max ECL Western Blotting Substrate (Bio-Rad laboratories, 1705062) was used with a ChemiDoc MP Imager System (Bio-Rad Laboratories).

### Immunofluorescence and cell staining

Cells were grown to desired conditions in 4-well chambered cover glass (Cellvis, C4-1.5H-N) or in Corning black 96-well clear bottom polystyrene microplates (Milliopore Sigma, CLS3904). Growth media was removed, and monolayers were washed with PBS (AmericanBio, Inc) prior to fixation with 4% PFA solution (Santa Cruz Biotechnology, Inc) for 20 min at room temperature. Fixed monolayers were washed three times, then incubated for 1 h in permeabilization buffer (PBS, 1% BSA, 0.1% saponin) at room temperature with rocking. The primary antibody was diluted in fresh permeabilization buffer based on manufacturers recommendation and cells were incubated overnight at 4°C with rocking. Cells were washed (10 min) three times with PBS, the secondary antibody was added in fresh permeabilization buffer (1:2000) for 2 h at room temperature in the dark with rocking. Cells were washed (10 min) three times with PBS and imaged within 24 h. To stain the cell nucleus, DAPI (Thermo Fisher Scientific, 62247) was added to the first wash of the final set of PBS washes. To stain bacterial nucleic acids, Hoechst 33342 (Thermo Fisher Scientific, H3570) was added along with secondary antibodies. F-actin was stained with Phalloidin DyLight 650 (Thermo Fisher Scientific, 21838) during the incubation with secondary antibodies.

Dilutions of primary antibodies were as follows: mouse monoclonal anti-ARP3 [FMS338] (Millipore Sigma, A5979), 1:500; mouse monoclonal anti-LAMP1 [H4A3] (Abcam, ab25630), 1:100; rabbit monoclonal anti-Paxillin [Y113] (Abcam, ab32084), 1:250; rabbit polyclonal anti-VASP (Millipore Sigma, HPA005724), 1:400; mouse monoclonal anti-Vinculin [7F9] (Santa Cruz Biotechnology, Inc; sc-73614), 1:250; mouse monoclonal anti-Xinα [D-8] (Santa Cruz Biotechnology, Inc; sc-166658) 1:200; rabbit polyclonal anti-XIRP1 (Thermo Fisher Scientific, PA5-53487), 1:1000; rabbit polyclonal anti-XIRP1 (Thermo Fisher Scientific, PA5-48605), 1:250. Secondary antibodies from Thermo Fisher Scientific were as follows: Alexa Fluor 488 donkey anti-mouse (A21202); Alexa Fluor 488 goat anti-rabbit (A11034); Alexa Fluor 568 goat anti-mouse (A11004); Alexa Fluor 568 donkey anti-rabbit (A10042).

### Microscopy, image processing, and fluorescence measurements

Samples in clear-bottom 96-well plates were imaged using an ImageXpress Pico Automated Cell Imaging System (Molecular Devices). Samples in 4-well chambered cover glass were imaged with either a Nikon epifluorescence microscope, Nikon TiE inverted spinning disk confocal, or a DeltaVision OMX SR Imaging System (GE Healthcare). Reconstruction of super-resolution images was done with softWoRx (GE Healthcare). Live-cell microscopy images were acquired using an environmental chamber (37°C, 5% CO_2_) with the widefield mode of the DeltaVision OMX SR Imaging System. Live-cell images were deconvoluted with softWoRx. Adjustments to image brightness and contrast was applied to entire images using Fiji/ImageJ 1.53c (Wayne Rasband, National Institutes of Health, USA).

The Fiji-based macro Poji (Herzog et al., 2020) was used to measure fluorescence intensity of individual podosomes and to determine podosome number per cell. Confocal microscopy was used to acquire Z-stacks at the base of podosomes marked by vinculin. For consistency, the Z-slice with the highest vinculin signal was selected for measurements associated with phalloidin staining of F-actin.

### RNA-Seq analysis and data mining

Hs27 cells were cultured in DMEM (10% FBS, 2 mM glutamine) to ∼70-80% confluency in T-175 flasks. Duplicate samples were either left untreated or treated with 50 ng/ml IFN-γ for 24 h. Cells were harvested using 0.25% trypsin/EDTA (Thermo Fisher Scientific, 25200056), and RNA was isolated using the Direct-zol RNA miniprep kit (Zymo Research, R2051). Total RNA was sent to GENEWIZ for standard library preparation, PolyA selection, and RNA sequencing (Illumina HiSeq, 2 x 150 bp).

To mine macrophages RNA-seq datasets, an R-script downloaded from the ARCHS^4^ website was used to generate a matrix containing gene-level counts from 1,933 macrophage datasets available from the Sequence Read Archive (SRA) (Lachmann et al., 2018). Quantile normalization was performed using Bioconductor (biocLite) and the preprocessCore package. Normalized XIRP1 expression was plotted with GraphPad Prism and M1-polarized THP-1 macrophage datasets (GEO accession: GSM3729256-8) were identified based on high XIRP1 expression (top 1.5% of all datasets). Raw fastq files from the SRA were downloaded along with files from control THP-1 monocyte samples (GEO accession: GSM3729250-2).

The Farnam computer cluster from the Yale Center for Research Computing was used for RNA-seq data analysis following a pipeline from rnabio.org (Griffith et al., 2015). Briefly, sequencing adapters were trimmed with Flexbar and quality of reads evaluated with FASTQC; reads were aligned using HISAT2 and read counts generated with HTSEQ; differential expression analysis and statistical significance was performed with edgeR using the exact-test and adjusting *p-values* based on a 5% false discovery rate. Volcano plots of DEGs were generated with GraphPad Prism. Plots exclusively containing actin-associated genes were filtered with the BioVenn web application based on Gene Ontology Terms (GO:0015629, GO:0003779, GO:0005884, GO:0031941, GO:0045010, GO:0007014).

### Phylogenetics and other bioinformatic tools

An alignment containing amino acid sequences from 221 XIRP1 homologues was generated using the online *MAFFT* FFT-NS-i algorithm (mafft.cbrc.jp/alignment/server). A maximum likelihood tree with a JTT matrix-based model was constructed using *MEGA* version 10.1.8 (Jones et al., 1992; Kumar et al., 2018). For clarity, tree branches were collapsed among closely related species. Protein domains were analyzed with *NCBI Orthologs*, the *Conserved Domain Database*, and the *Conserved Domains-Search* tool (NCBI). Gene synteny analysis within the +/-1,000,000 base pair region flanking XIRP1 was performed with the NCBI Genomic Context browser. Immunity-related genes (*e.g.*, MYD88, CX3CR1, CCR8) were identified within the human XIRP1 region and presence or absence of genes was determined in representative organisms from mammals (*P. troglodytes, M. musculus, O. anatinus*), birds (*D. novaehollandiae*), reptiles (*A. mississippiensis*), amphibians (*M. unicolor, X. laevis*), fish (*D. rerio, L. oculatus, C. milii*), and sea lamprey (*P. marinus*).

Predicted binding sites for transcription factors were identified through the *TF Analysis* online tool (interferome.org). Independent verification was done through the Swiss Institute of Bioinformatics *Eukaryotic Promoter Database* (epd.epfl.ch).

### Experimental animals and mouse infections

Animal experiments were approved by the Yale University Institutional Animal Care and Use Committee (Protocol #2020-11647). Investigators were not blinded and animal studies were not randomized. Animal studies were gender-matched and mice were 8-13 weeks old. C57BL/6NJ mice were purchased from Jackson Laboratories (Bar Harbor, ME) and maintained under specific pathogen-free conditions at the Yale West Campus Animal Research Facility. XIRP1-knockout mice were obtained from the laboratory of Da-Zhi Wang at Harvard University (Gustafson-Wagner et al., 2007); breeding, genotyping, and husbandry was performed at Yale University.

*Listeria* cultures were grown to OD_600_ = 0.6 as previously indicated. Bacteria were centrifuged (10,000 x g, 3.5 min), washed and resuspended in DPBS. Mice were infected intraperitoneally with an inoculum of 7.5×10^5^ CFU per mouse in a 200 μl volume. Mice were monitored at least once daily for fourteen days and euthanized upon reaching a moribund state characterized by hunched posture, immobility, and a drop in abdominal surface temperature bellow 27°C. To determine bacterial burden, mice were euthanized four days post-infection, organs were harvested and homogenized in DPBS. Serial dilutions of homogenates were plated in BHI agar plates containing 200 μg/ml streptomycin.

## Supporting information

Supplemental Table 1 Fibroblast DEGs

Supplemental Table 2 Macrophage DEGs

## Acknowledgments

The authors thank Joerg Nikolaus (Yale West Campus Imaging Core) for technical help and advice with confocal microscopy.

## Additional information

### Funding

| 645 | **Funder** | **Grant reference number** | **Author** |
| --- | --- | --- | --- |
| 646 | HHMI | GT11834 | Rodolfo Urbano |
| 647 | NIH | 2 R01 AI108834-07 | John D. MacMicking |
| 648 | NIH | 3 R01 AI068041-14 | John D. MacMicking |

John D. MacMicking is an Investigator of the Howard Hughes Medical Institute.

## Author contributions

Rodolfo Urbano, Conceptualization, Data curation, Formal analysis, Funding acquisition, Investigation, Methodology, Visualization, Validation, Writing – original draft, Writing – review and editing; Eui-Soon Park, Kyle Tretina, Alexandru Tunaru, Ryan G. Gaudet, Xiaoyun Hu, Da-Zhi Wang, Methodology, Resources; John MacMicking, Conceptualization, Funding acquisition, Project administration, Supervision, Writing – review and editing.

## Author ORCIDs

Rodolfo Urbano https://orcid.org/0000-0001-7404-774X Eui-Soon Park https://orcid.org/0000-0002-1375-8027

Kyle Tretina https://orcid.org/0000-0002-1740-2730 Ryan G. Gaudet https://orcid.org/0000-0001-8496-3653 John D. MacMicking https://orcid.org/0000-0002-1734-135X

## Additional files

### Supplementary files

- Table 1 Fibroblast DEGs
- Table 2 Macrophage DEGs

---

**Figure supplement 1.**
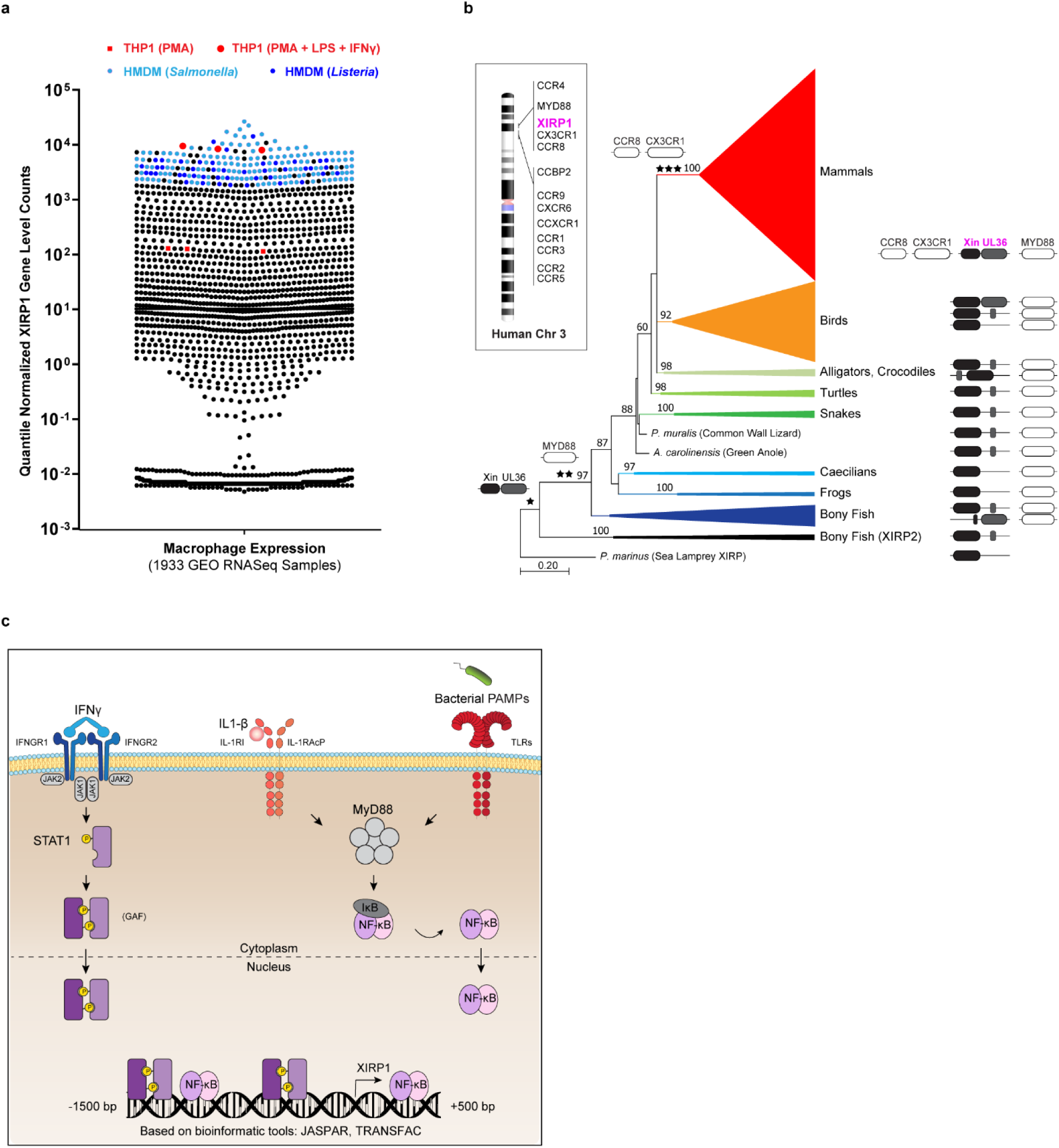
*In silico* analysis linking XIRP1 to host immune processes (a) XIRP1 gene-level counts across human macrophage RNA-Seq datasets obtained from ARCHS^4^. Samples from THP-1 derived macrophages are shown in red and from human monocyte derived macrophages are shown in blue (*Salmonella-*infected, light blue; *Listeria*-infected, dark blue). (b) Maximum-likelihood tree with JTT matrix-based model shows the distribution of XIRP1 across vertebrates. Shown are full or partial Xin and UL36 domains identified through the conserved domain database. The presence of *MYD88* and genes encoding macrophage chemokine receptors within a +/-1,000,000 base pair region flanking XIRP1 is indicated. Additional genes within two chemokine receptor clusters on human chromosome 3 are shown. (c) Model shows immune pathways that stimulate XIRP1 expression and putative transcription factor binding-sites on the XIRP1 regulatory region (bioinformatically predicted with indicated webtools).

**Figure supplement 2.**
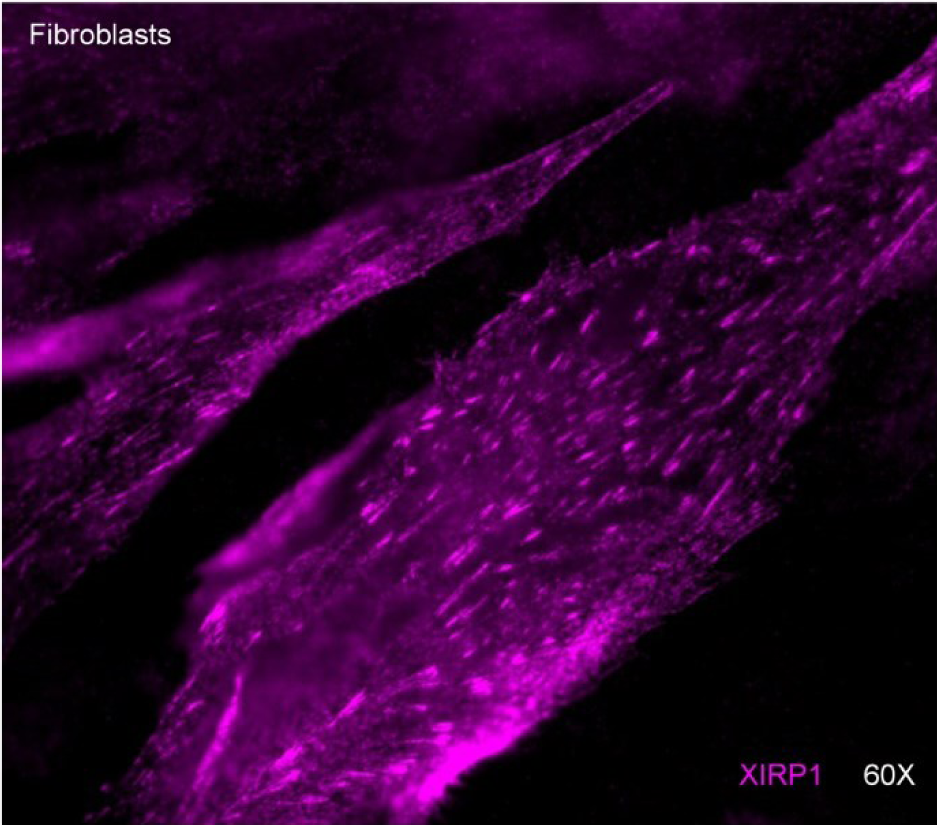
Immunostaining of Hs27 fibroblast showing XIRP1 localization to focal adhesions at the base of the cell.

